# Performance - Based Clustering Enables the Study of Physiological Features Supporting Sustained Attention Capacities

**DOI:** 10.1101/497206

**Authors:** José Eduardo Marques-Carneiro, Bérengère Staub, Anne Bonnefond

## Abstract

In the present study, we investigated different sustained attention performance profiles in the general adult population and if age is a determinant factor among these profiles. We also reported some specificities in terms of brain oscillations. The sustained attention to response task was applied combined with electrophysiological recordings in 59 adults, aged from 19 to 86 years. We used a hierarchical cluster analysis to group individuals based on similarity of levels and patterns of performance across variables of interest. We focused on the most common attentional–related variables, which allowed us to identify four distinct clusters showing specific sustained attention profiles. Furthermore, our analysis clearly revealed that age was not the determinant factor of these profiles. Finally, we highlighted specificities in terms of brain oscillations. Subjects from cluster 1– the “high-performers” – demonstrated a globally increased theta-gamma coupling maintained over time and an increase of theta activity at the FC6 and F8 electrodes. Subjects from cluster 2, who adopted a more cautious response strategy, presented a higher level of modulation exerted by the electrode FT7 over Oz and a higher theta-gamma coupling at the Oz. This result suggests that cluster-based method to useful in understanding specific mechanisms underlying sustained attention features.

## 1. Introduction

Sustained attention, also called vigilance or vigilant attention, is defined as the ability to efficiently sustain conscious stimulus processing during long periods of time (more than a few seconds) (Parasuraman et al. 1989). This “stimulus processing” refers to the simple detection and/or discrimination of stimuli, including simple cognitive or motoric responses, but excluding “higher” attentional or executive functions (e.g. dividing attention, spatial orienting, resolving interference, or selecting between several overt responses) (Langner and Eickhoff 2013). The assessment of sustained attention ability is achieved through different measures of performance reflecting either attentional fluctuations (Macdonald et al. 2011; Esterman et al. 2013) or attentional deteriorations (Robertson et al. 1997; Wascher et al. 2014). The most widely sustained attention task is the sustained attention to response task (SART (Robertson et al. 1997), in which subjects are tasked with withholding responses to a predefined rare numeric target (e.g., 3) and overtly responding to a larger digit set (e.g., 1, 2, 5, 7, 6, 7, 8, and 9). In this Go/No-Go task, the classical measures of performance used are the number of commission errors (CE), the median (or mean) reaction time (RT) but also the moment-to-moment variability of RT, such as RT standard deviation (RT SD). RT variability has become a widely used dependent measure in the SART literature as it is believed to reflect subtle differences in RT that are produced by lapsing attention (Allan Cheyne et al. 2009; Seli et al. 2012). Slope variability, which refers to the change of RT over the course of the task, is less frequently used. However, the subject’s tendency to slow down or to speed up as the task progresses is also highly informative about their ability to sustain attention over time (Lundwall and Watkins 2015). A slow-down over the course of the task associated with high rates of errors may reflect difficulty in maintaining attention over time, whereas when associated with a small number of errors, it may rather reflect the adoption of a more cautious response strategy in order to avoid errors. To our knowledge, such differences in speed-accuracy trade-off have been evidenced only by comparing different age groups. Indeed, in comparison to young people, the elderly commit less CE and tend to slow down throughout the task (Brache et al. 2010; Carriere et al. 2010; Jackson and Balota 2012; Staub et al. 2014). However, in these studies, it is possible that because subjects have been examined as different age groups (younger versus elderly), important individual differences that could categorize participants into sustained attention performance profiles subgroups have been missed. The identification of such profiles could help increase understanding of fundamental mechanisms underlying sustained attention ability.

In light of this, the primary aim of the present study was to determine whether sustained attention performance profiles exist in the general adult population. Data-driven analyses are commonly used to identify cognitive profiles (Miller et al. 2012; Liepelt-Scarfone et al. 2012; Hawkins et al. 2015; Reser et al. 2015). Here, we used a cluster analysis, which groups individuals in terms of similar levels and patterns of performance across variables of interest (Clatworthy et al. 2005). We focused on the four key variables described above (mean number of CE, median RT, RT SD and slope variability). A second aim was to examine potential age differences among these performance profiles. The subjects involved in our study were aged between 19 and 86 years. Finally, in an attempt to explain the presence of different performance profiles, we examined the specificities in term of oscillatory activity over time across the profiles. To that end, we used an oscillatory model of sustained attention recently proposed by Clayton et al (Clayton et al. 2015). Within this framework, sustained attention relies on (1) cognitive monitoring and evaluation of ongoing cognitive processes mediated by frontomedial theta oscillations (**Figure 1A**) and (2) selective excitation of task-relevant cortical areas through gamma oscillations (**Figure 1B**).

**Figure 1:**
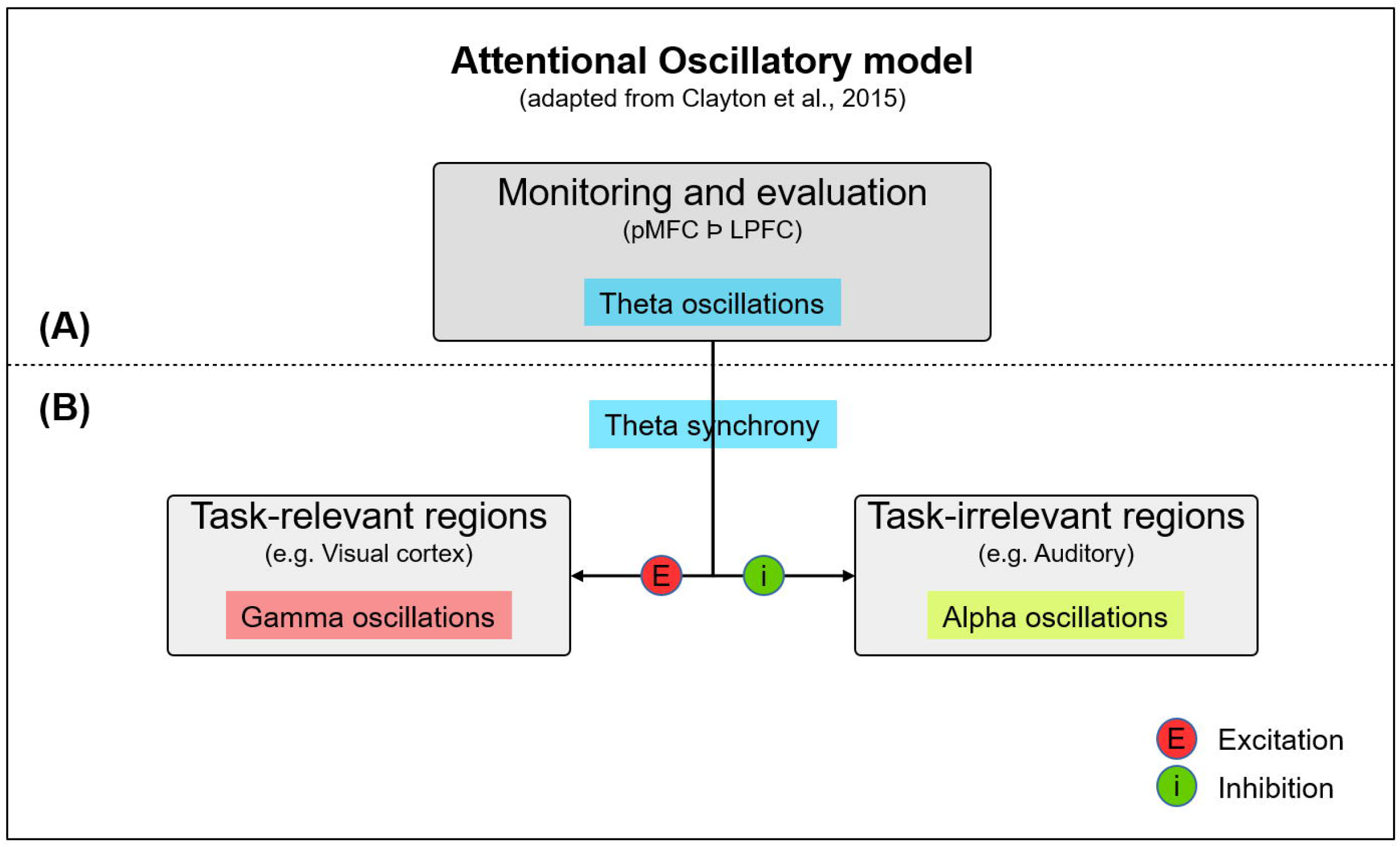
Attentional oscillatory model (adapted from Clayton et al., 2015). **(A)** monitoring and evaluation of relevant activity supported by theta oscillations in fontal cortices and by communication between posterior medial frontal cortex (pMFC) and lateral prefrontal cortex (LPFC). **(B)** Synchronization at theta band between frontal regions and taks-relevant and task-irrelevant regions.

## 2. Methods

All experiments were approved by the “Comité de Protection des Personnes” (CPP) Est-II and were performed in accordance with relevant guidelines and regulations.

### 2.1 Subjects

Fifty-nine subjects (28 males; mean age: 52.6 years; range: 19-86 participated in this experiment. The level of education was 14 (± 3) years. All subjects declared that they were free of neurological and psychiatric diseases and had normal or corrected-to-normal vision. All subjects gave their written informed consent.

### 2.2 Task and performance measures of sustained attention ability

All participants completed a 30-min task – the sustained attention to response task (SART; a Go/No-Go task in which digits ranging from “1” to “9” were presented in a random order. Subjects were instructed to respond as quickly and accurately as possible to the digits with a press on the control key of the keyboard upon presentation of each digit with the exception of the digit 3, which required response inhibition. Each digit was presented for 150 ms followed by an inter-stimulus interval (ISI) that varied randomly between 1,500 and 2,500 ms. All digits, including the 3, were presented with equal probability. Five randomly allocated digit sizes were presented to increase the demand for processing the numerical value and to minimize the possibility that subjects would set a search template for some perceptual feature of the target digit (“3”). The digit font sizes were 100, 120, 140, 160 and 180 in Arial font. The five allocated digit sizes subtended vertical angles of 1.39°, 1.66°, 1.92°, 2.18° and 2.45°, respectively, at a viewing distance of 70 cm. The digits were presented in black, 0.25° above a central yellow fixation cross on a grey background, on a standard 17-inch computer screen. Ninety consecutive blocks of trials were used, each block corresponding to a random presentation of all nine digits.

Four dependent variables were examined. The variables are: (1) the total number of commission errors (CE); (2) the median reaction time (RT); (3) the variability of reaction time (RT SD); and (4) the slope variability (slope). For the slope calculation, we used the sample Pearson correlation coefficient to access the linear correlation between the median RT and the block number (1 to 90). A positive coefficient suggests a slowing down over the course of the task, a negative coefficient points towards a speeding up.

### 2.3 EEG recordings and analysis

Electroencephalograms (EEGs) were recorded from 64 Ag/Acl BioSemi active electrodes mounted in an elastic cap. Electrodes were placed according to the standard 10–20 system designed to cover a large median area of the scalp. An online band-pass filter was applied (0,01 – 100 Hz). Data were sampled at a rate of 512 Hz. For EEG analysis, the task was divided into six periods (p1/p2/p3/p4/p5/p6), with each period corresponding, in terms of duration, to 5 mins.

#### Power spectrum density (PSD)

To evaluate the influence of theta oscillations during the sustained attention task we extracted PSD using the Welch method, using a window length of 1 sec and a window overlap ratio of 50 %. The theta frequency was considered between 4-8Hz. The theta PSD was then calculated for the followings electrodes: Fz and Afz for posterior medial frontal cortex and FC5, FC6, FT7, FT8, F7 and F8 for lateral prefrontal cortex. PSD was performed with Brainstorm (Tadel et al. 2011), which is documented and freely available for download online under the GNU general public license (http://neuroimage.usc.edu/brainstorm).

#### Granger causality analysis

To evaluate the synchronism at theta-band frequency between regions considered in the oscillatory model, we performed a Granger Causality analysis. This methodology is used as a statistical hypothesis test for determining whether one time series may be useful in forecasting another, and therefore enabling analysis of the connectivity between two signals (Granger 1969). In summary, this analysis is based on vector autoregressive models to evaluate the causal relationship between two time series data such EEG. Granger causality analysis was also performed with Brainstorm (Tadel et al. 2011). We used a maximum Granger model order of 10, a maximum frequency resolution of 1Hz and the highest frequency of interest of 8 Hz. We calculated the granger causality to evaluate the modulation of Fz and AFz over the electrodes FC5, CF6, FT7, FT8, F7 and F8 as well as the modulation of FC5, CF6, FT7, FT8, F7 and F8 over occipital and parieto-occipital electrodes (Oz, POz, PO3 and PO4) and auditory regions (C5 and C6).

#### Phase-amplitude coupling (PAC)

To evaluate the modulation of task-relevant regions we used the PAC method. In the PAC (or nested oscillations) the low-frequency (nesting) phase will drive the power of a coupled higher frequency (nested). Therefore, we analyze the synchronization of amplitude envelope of gamma frequency with the phase of theta band. For the PAC analysis, region of interest was defined thanks to the granger causality analysis, only occipital regions modulated in the theta band were considered. PAC analysis was also performed with Brainstorm software (Tadel et al. 2011).

### 2.4 Statistical analysis

#### Aim 1

A cluster analysis, utilizing the four measures of performance (CE, RT, RT SD and slope), was used to identify sustained attention performance profiles. We used the Ward Hierarchical Clustering (hclust function in R) to generate clusters based on this performance measures. The minimal sample size suggested for hierarchical cluster analysis is of 2^k^ (k=number of variables) (Formann 1984). Consequently, as we have four variables in this study, we have more than three times the suggested minimal sample size. A dendogram was made to plot the clusters and a radial plot was used to represent graphically the attentional performance profiles of the identified cluster groups after standardizing all values with a z-score. Measures of performance (CE, RT, RT SD and slope) were subjected to a one-way ANOVA, which included the between-subject variable Cluster.

#### Aim 2

Age was subjected to a one-way ANOVA, which included the between-subject variable cluster. To explore further we also performed an analysis of covariance (ANCOVA) in order to control for a possible confounding aging effect.

#### Aim 3

Data from the PSD, Granger causality and PAC, were subjected to a two-way ANOVA including the between-subject factor Cluster and the within subject factor Period (p1/p2/p3/p4/p5/p6).

For all statistical analysis (ANOVA and post-hoc test) we considered a probability lower than 5 % (p < 0.05) to commit a type I error. For all analyses, when significant differences were found, the post-hoc test Bonferroni was used considering all main effects and interactions. A multiple comparison correction (familywise error rate) was applied for each statistical analysis (FWE ≤ 1 – (1 − αIT)^c^; where αIT = alpha level for an individual test (e.g. .05) and c = Number of comparisons). All statistical analyses were performed using the Statistica software.

## 3. Results

### 3.1 Aim 1: Identification of sustained attention performance profiles

Ward Hierarchical Clustering method generated 4 clusters. A dendogram shows the distribution of each cluster (**Figure 2**).

**Figure 2:**
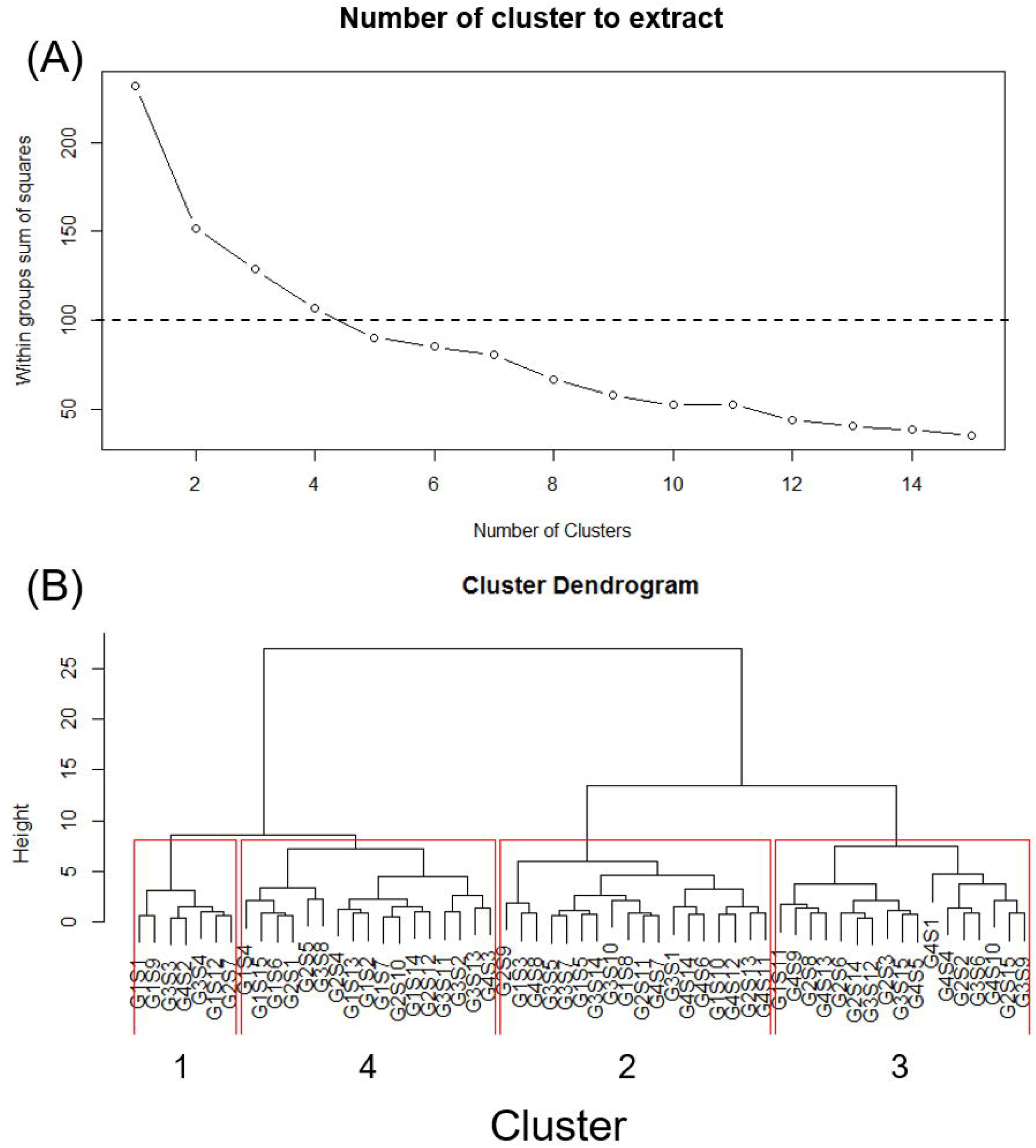
Cluster extraction. **(A)** Four clusters is the optimal number of cluster to extract; **(B)** Cluster dendrogram.

Cluster 1 includes 7 subjects with a low number of CE, a short RT, a low RT SD and a low slope. In cluster 2, we found 18 subjects with a high slope (slow down), medium RT and RT SD and a medium number of CE. Cluster 3 included 17 subjects with a long RT, a high RT SD, a low number of CE and a low slope. Finally, cluster 4 included 17 subjects with a short RT, a low RT SD, a high number of CE and a low slope (**Table 1**). Correlations between performance variables are shown in supplementary **Figure S1**.

**Table 1:** Cluster’s performance: CE, RT, RT-SD and Slope (mean ± SD).

|  | CLUSTER 1 | CLUSTER 2 | CLUSTER 3 | CLUSTER 4 |
| --- | --- | --- | --- | --- |
| CE | Low<br>(16 ± 3.05) | Low<br>(19.33 ± 10.83) | Low<br>(11.29 ± 6.10) | High<br>(39.05 ± 8.90) |
| RT | Short<br>(297.17 ± 36.25) | Medium<br>(348.36 ± 50.28) | Long<br>(396.44 ± 63.98) | Short<br>(279.76 ± 49.78) |
| RT SD | Low<br>(37.18 ± 7.63) | Medium<br>(56.91 ± 11.56) | High<br>(71.20 ± 17.29) | Low<br>(41.98 ± 12.72) |
| Slope | Low<br>(0.07 ± 0.16) | High<br>(0.45 ± 0.14) | Low<br>(0.05 ± 0.24) | Low<br>(0.01 ± 0.28) |

RT – The ANOVA revealed the following main effect of cluster (F(3,55)= 15.175; p<0.001; ɳ^2^= 0.45; power=0.99). The post-hoc test revealed that RT in cluster 3 is longer than in clusters 1 and 4, and shorter in cluster 4 than in cluster 2 (**Figure 3A**).

CE – The ANOVA revealed a main effect of cluster (F(3,55)= 34.031; p<0.001; ɳ^2^= 0.64; power=1). The post-hoc test revealed that the mean number of CE in cluster 4 is higher than in the three other clusters and lower in cluster 3 than in cluster 2 (**Figure 3B**).

RT SD – The ANOVA revealed a main effect of cluster (F(3,55)= 17.624; p<0.001; ɳ^2^= 0.49; power=0.99). The post-hoc test revealed that RT SD in cluster 3 is higher than in the three other clusters. The RT SD in cluster 2 is also higher than in clusters 1 and 4 (**Figure 3C**).

Slope – The ANOVA revealed a main effect of cluster (F(3,55)= 13.780; p<0.001; ɳ^2^= 0.42; power=0.99). The post-hoc test revealed that slope in cluster 2 is higher than in the three other clusters (**Figure 3D**).

**Figure 3:**
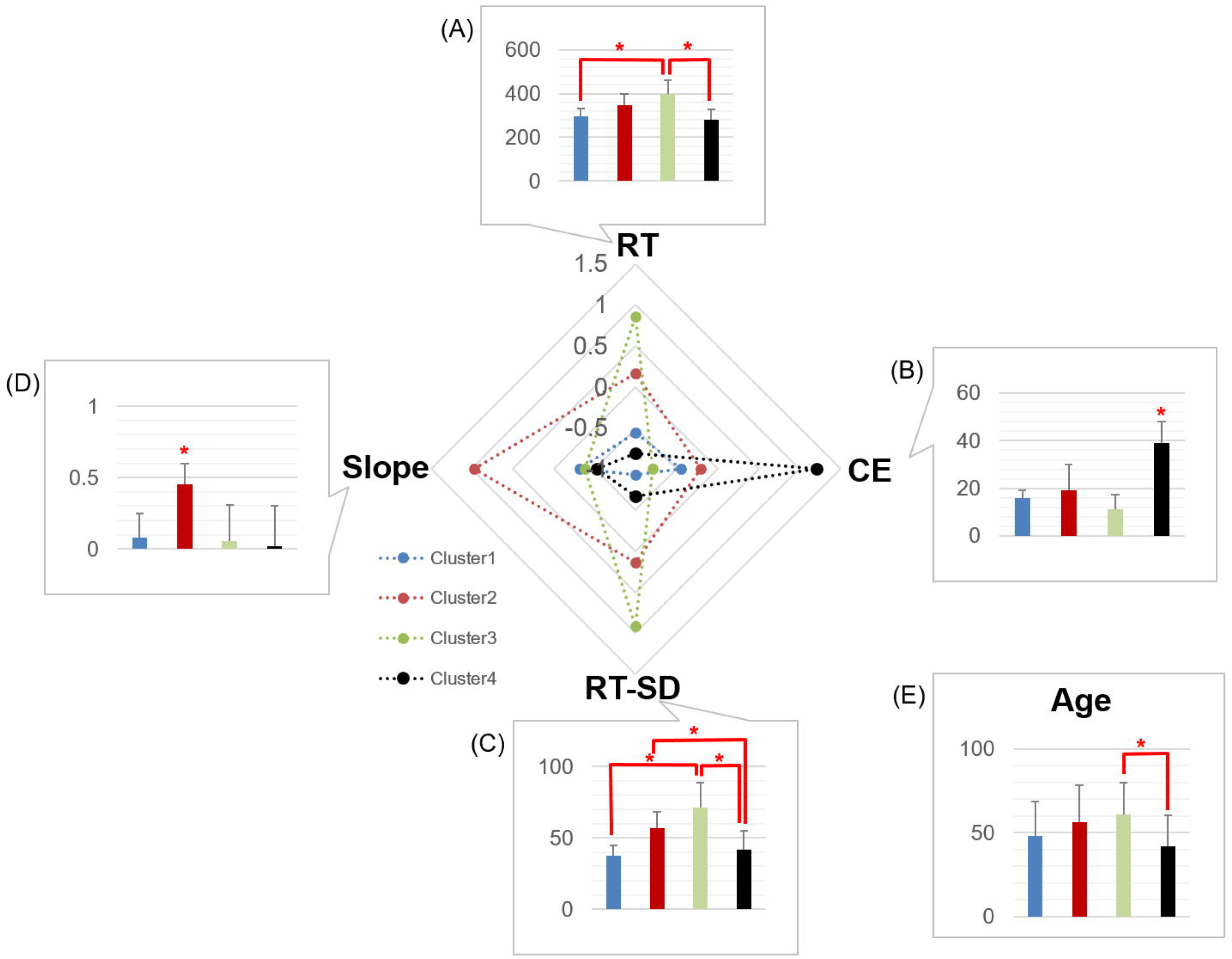
Performance profile. **(A)** Reaction time – RT; **(B)** Commission error - CE; **(C)** reaction time standard deviation – RT-SD; **(D)** Reaction time slope - Slope; **(E)** Age. All values are presented as mean and ± standard deviation.

### 3.2 Aim 2: Investigation of age effects

Age – The age distribution among the four clusters is shown in the supplementary figure S2. The ANOVA revealed a main effect of cluster (F(3,55)= 2.896; p=0.043; ɳ^2^= 0.13; power=0.65). Post-hoc tests revealed a lower age of cluster 4 (43.53 ±17.83) compared to cluster 3 (59.74 ±19.43), while no difference was observed between clusters 1 (46.37 ±23.16) and 2 (56.84 ±21.99) (**Figure 3E**).

The ANCOVA made on each dependent variable with Age as covariant showed the same cluster effect (**Table 2**). Post-hoc tests revealed also the same differences as those described above.

**Table 2:**
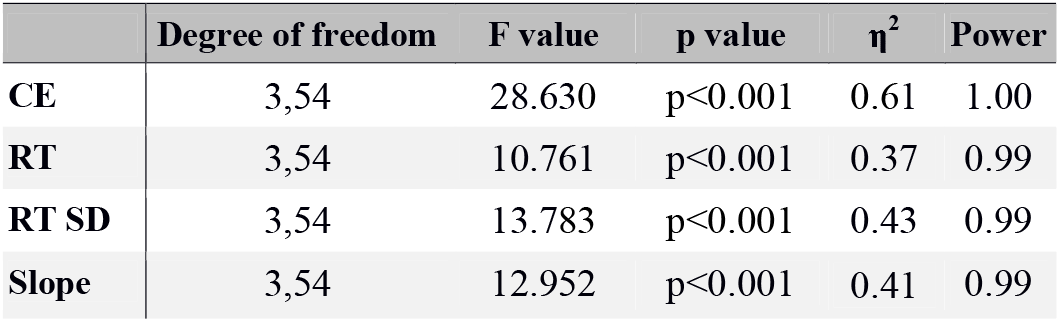
Results of the ANCOVA with Age as covariant.

### 3.3 Aim 3: Investigation of profile’s specificities in terms of oscillatory activity over time

To verify if Clayton’s model (Clayton et al. 2015) may explain the differences observed between clusters, we focused on two main processes of this model: the monitoring and activation of relevant structures. Complete result table is available in supplemental material, only results from the electrodes highlighting statistical differences are presented here.

Monitoring - First we evaluated the influence of theta power on electrodes located on posterior medial frontal cortex (Fz and AFz) and lateral prefrontal cortex (FC5, FC6, FT7, FT8, F7 and F8). The ANOVA performed on theta power revealed an interaction between period and cluster in two of the 8 electrodes analyzed: FC6 (F(15,275)= 2.325; p=0.023; ɳ^2^= 0.11; power=0.98) and F8 (F(15,275)= 2.232; p=0.039; ɳ^2^= 0.10; power=0.97). The post-hoc test revealed that theta power was significantly different over periods in cluster 1, in which theta power was higher in periods 5 and 6 than in periods 1 and 2 at FC6 (**Figure 4A**) and at F8 (**Figure 4B**). It remained stable all along the task in the three other clusters.

**Figure 4:**
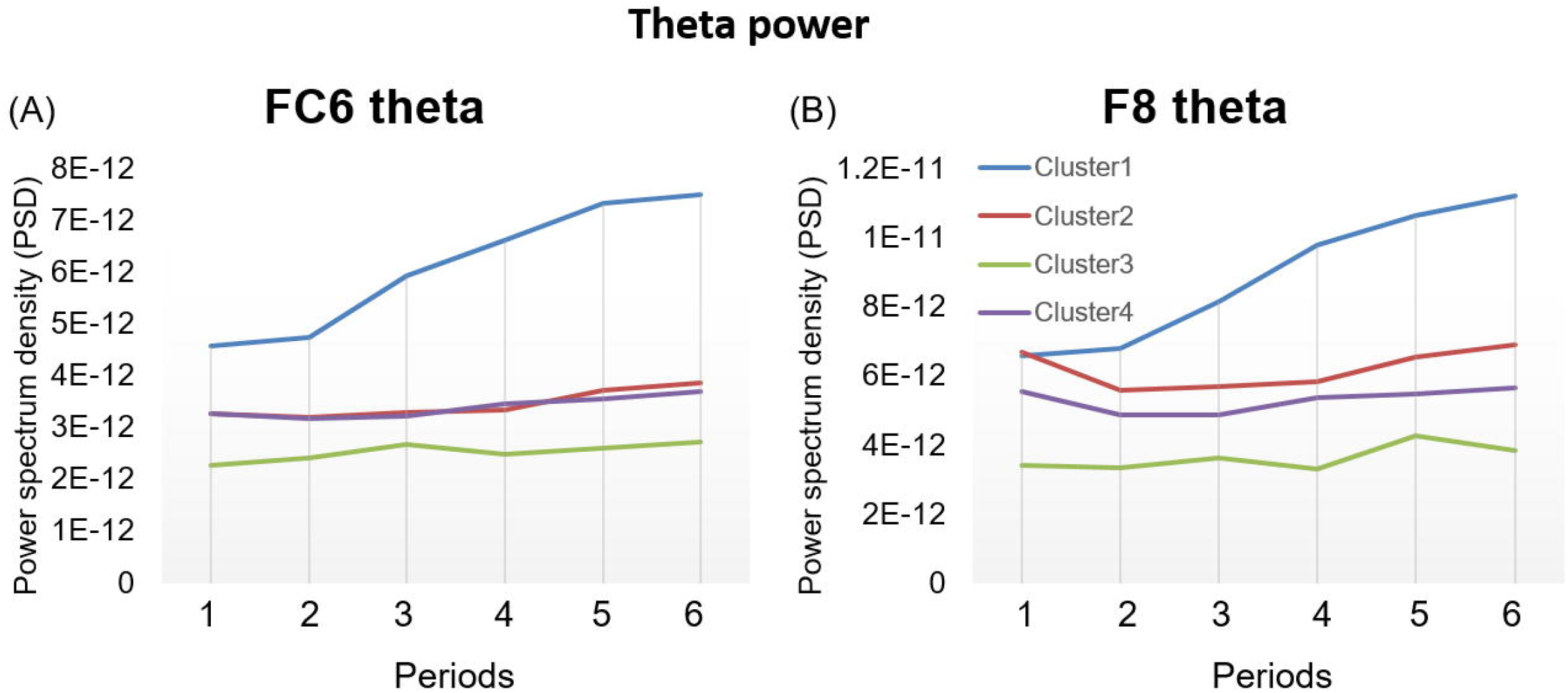
Theta power in frontal regions. **(A)** Theta power in fronto-central cortex - FC6; **(B)** Theta power in frontal cortex – F8.

Activation of task-relevant region – We used a granger causality analysis at theta band to evaluate the relationship between monitoring regions (posterior medial frontal and lateral prefrontal cortices) and visual area. The ANOVA of the Granger causality value at theta-band revealed an interaction between period and cluster for the causal connectivity between the electrodes FT7 and Oz (F(15,275)= 2.628; p=0.023; ɳ^2^= 0.12; power=0.99). The post-hoc test revealed that cluster 2 showed a higher theta connectivity in period 1 compared to periods 5 and 6 (**Figure 5A**). In addition, we observed a negative correlation of −0.40 between FT7-Oz theta connectivity and RT over the 6 periods, while we found a correlation of 0.22, 0.14 and −0.03 for the clusters 1, 3 and 4, respectively. The **Figure 5B** shows the variation of RT over the different blocks.

**Figure 5:**
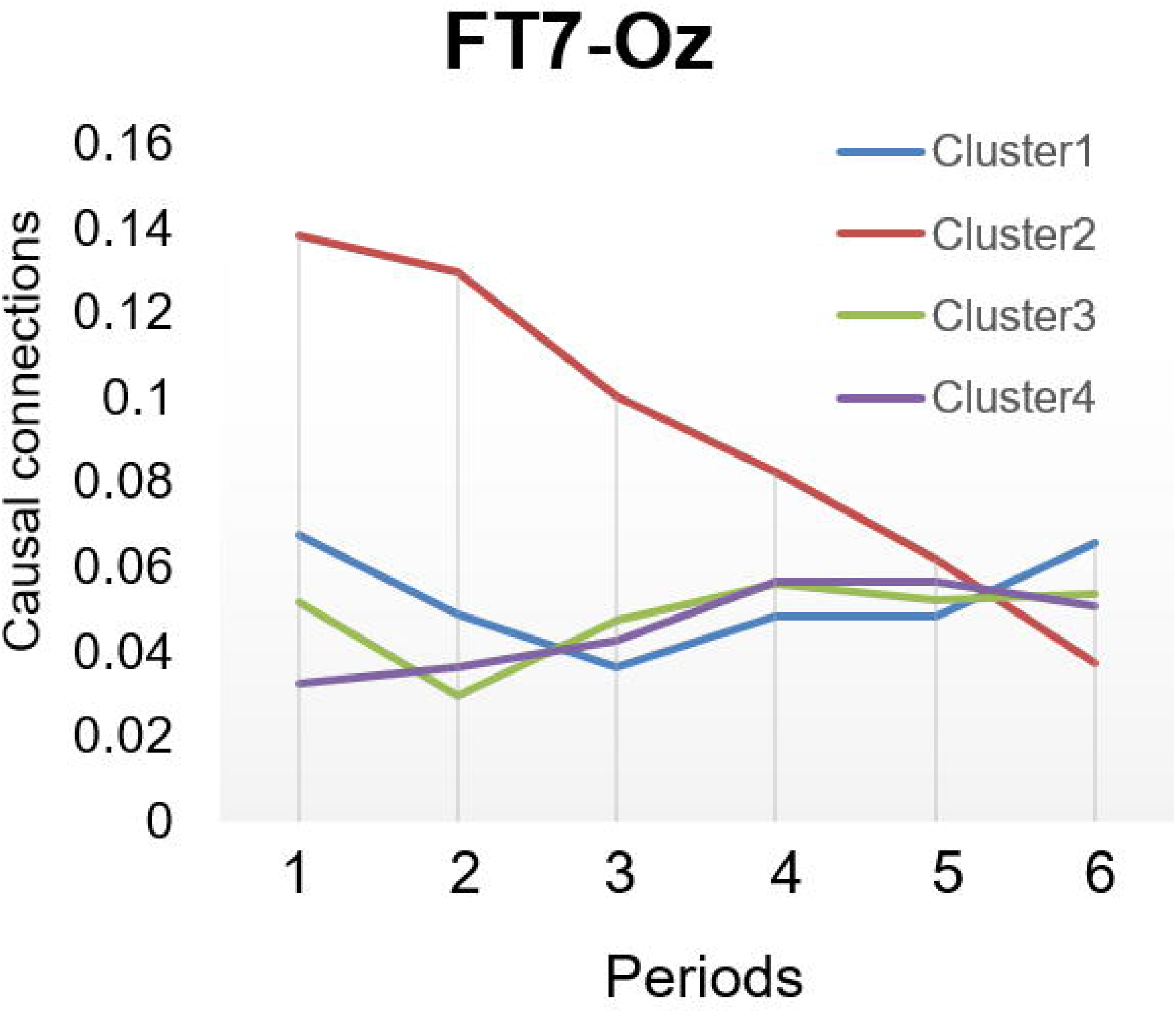
Activation of task relevant structures calculate with Granger causality connection between FT7 and Oz.

As Oz was found to be modulated by FT7 at theta band synchrony, we analyzed the theta-gamma coupling on this electrode. The **Figure 6** shows the theta-gamma coupling for the four clusters over time. Descriptively, in this figure we observe clear differences among the four clusters. We note that cluster 1 had a higher and over-time stable theta-gamma coupling in a wide-range of regions. We also observed that in the occipital region, the theta-gamma coupling was higher in the first period for the subjects of cluster 2, but decreased over time. The ANOVA for the theta-gamma coupling in occipital region showed an interaction between cluster and period in the Oz electrode (F(15,275)= 2.086; p=0.010; ɳ^2^= 0.10; power=0.96). The post-hoc test revealed that cluster 2 showed a lower theta-gamma coupling on the period 5 compared to the period 1 (**Figure 7**).

**Figure 6:**
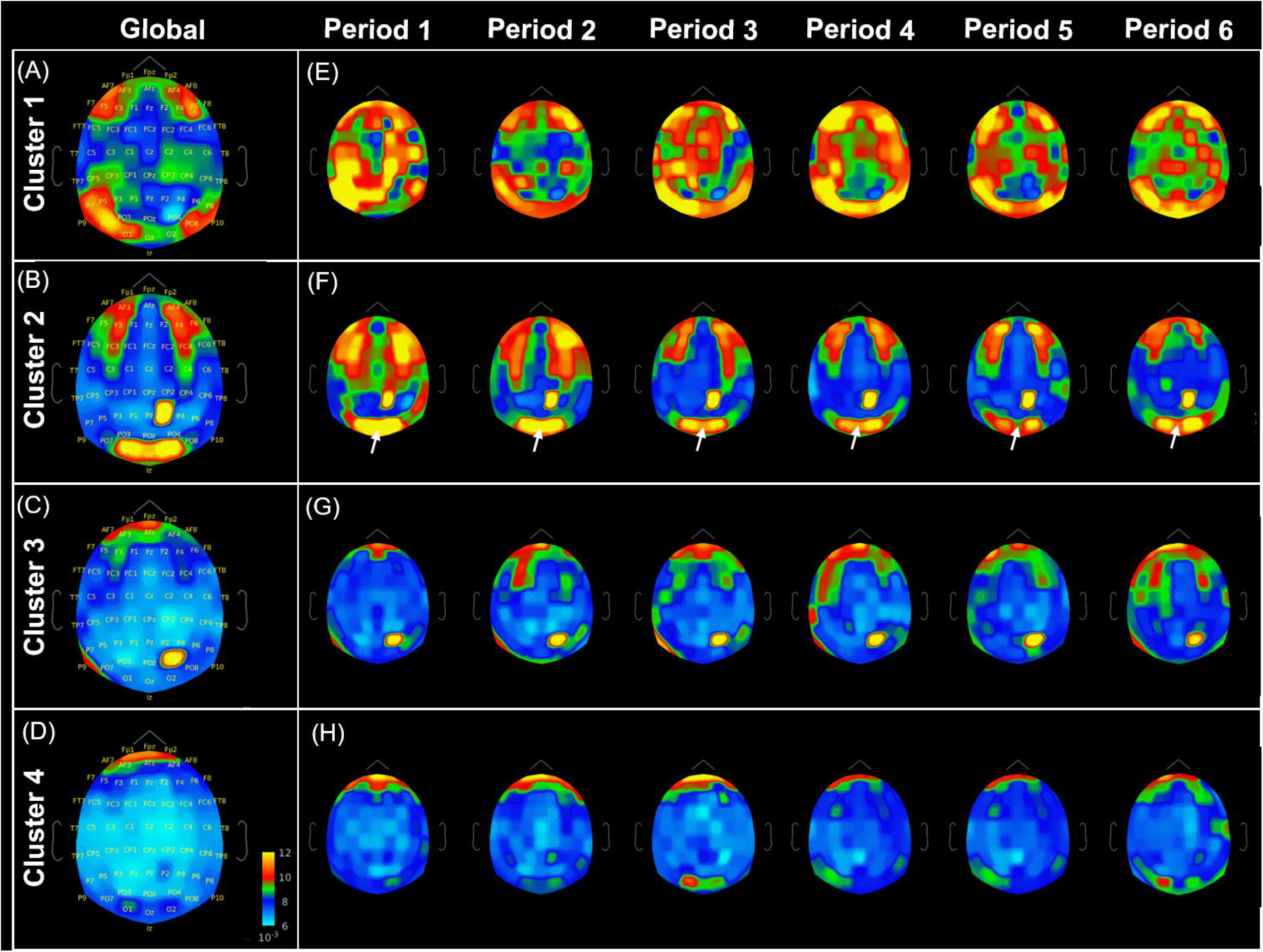
Cross-frequency coupling. **(A, B, C and D)** Global coupling between theta phase and gamma power. **(E, F, G and H)** Over time cross-frequency coupling. The arrows indicates a significant difference observed in occipital regions in cluster 2 subjects (see. Figure 7).

**Figure 7:**
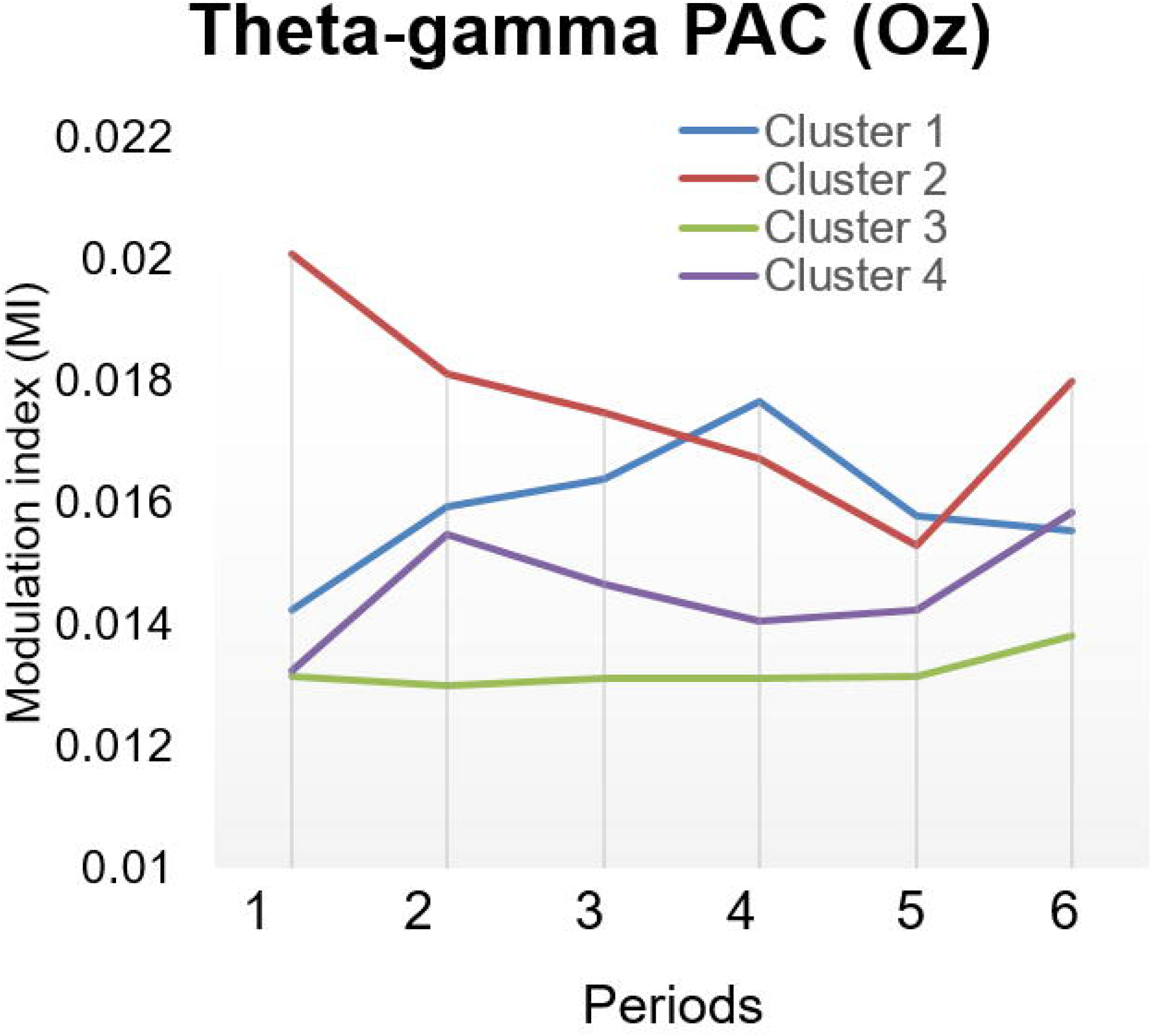
Modulation index of theta-gamma coupling in occipital cortex – Oz (Visual region).

## 4. Discussion

The main aim of the present study was to determine sustained attention performance profiles in the general adult population. To that end, we applied the SART to 59 adults, aged from 19 to 86 years, and focused on four classical measures of performance. In addition, we examined age differences among these performance profiles and specificities in terms of oscillatory activity over time.

### Identification of sustained attention performance profiles

By clustering subjects with similar performance we were able to identify four distinct clusters (**Figure 3**; **Table 1**). Each of them differed from the others on at least one measure of performance. More precisely, the subjects included in cluster 1 were “high-performers”: they are accurate (low level of CE) and fast, they have few attentional lapses (low RT SD) and no attentional decline over time (negative slope). Subjects included in cluster 2 demonstrated a higher positive slope, reflecting a markedly RT slowdown over the course of the task. In association with a medium level of CE, this may suggest the adoption of a more cautious response strategy. Cluster 3 grouped subjects with the higher RT variability (RT SD), believed to reflect subtle differences in RT that are produced by attentional lapses (Allan Cheyne et al. 2009; Seli et al. 2012). Finally cluster 4 included subjects with the highest number of CE.

Further, our results show that the clusters 3 and 4 seem to include two contrasting populations that reflect opposite speed-accuracy trade-off related profiles. On one hand cluster 3 includes individuals with a high level of accuracy (indicated by a high RT and SD and a low number of CE), on the other hand cluster 4 includes faster subjects (with a fast RT, low SD, and a high number of CE), suggesting an opposite speed-accuracy trade-off balance. The relationship between speed and accuracy seems to be a signature of the decision process used (Heitz 2014), for example “subjects either used a guessing strategy or a highly accurate controlled strategy” (Swensson 1972). This kind of differences in speed-accuracy trade-off was observed between different age groups, in which was found that elderly commit less CE and tend to slow down throughout the task if compared to young people (Brache et al. 2010; Carriere et al. 2010; Jackson and Balota 2012; Staub et al. 2014). However, in this study the difference was attributed to an age-related difference excluding important individual differences, such different possible strategies. In this sense, this methodology enables us to study more deeply the neural circuitry and other physiological aspects supporting each one of these characteristics.

### Investigation of age effects

The investigation of age differences among the four clusters revealed an age difference only between clusters 3 and 4. Cluster 3 included subjects older (around 60 years old) than cluster 4 (around 43.5 years old). However, age had no impact on the indicators analyzed. Indeed, we still found similar differences in RT, SD, Slope and CE among the four clusters after controlling by age. In this sense, age seems not be a determinant factor which might explain or indicate the preferential utilization of one or other strategy. Thus, it may explain why in the literature we find very heterogeneous results regarding the relationship between sustained attention capacities and aging. However, the identification of cluster groups composed of similar performance profiles may allow us to go further and identify/study biomarkers related to specific strategies (e.g. controlled or automatic) as well as aspects associated with attentional features such attentional lapses.

### Oscillatory model of sustained attention

To take our observation further, we tested the oscillatory model of sustained attention recently proposed by Clayton et al (Clayton et al. 2015). This model suggests that sustained attention relies on two main functions, (1) monitoring and evaluation of ongoing cognitive processes mediated by theta oscillations on posterior medial frontal cortex and lateral prefrontal cortex and (2) a selective excitation of task-relevant cortical areas through gamma oscillations and inhibition of task-irrelevant areas through alpha oscillations. In the present study, we focused on the monitoring process as well as in the activation of relevant structures.

Regarding the first function (1) we studied the effects of task engagement on theta oscillations in electrodes located in monitoring relevant structures. Our results revealed that the “high-performers” (individuals from cluster 1) presented an increase over time of theta activity in right-frontal regions (FC6 and F8), which is not observed in the other clusters (**Figure 4**). It suggests that over the time course of the task there is an increasing need of theta activity to maintain a good performance. The fact that theta oscillations, which are believed to support long-range transmission of information, were increased in these subjects may indicate an enhanced modulatory activity (Clayton et al. 2015). Thus, the over-time increase of theta-band activity observed in the right-frontal region of the “high-performers” corroborates the importance of this cortical region in the maintaining of attention (Posner and Petersen 1990; Pardo et al. 1991; Rueckert and Grafman 1996; Coull 1998).

Regarding the activation of task-relevant regions (2), we observed that individuals included in cluster 2 presented a high level of functional connectivity at theta-band between FT7 and Oz electrodes at the beginning of the task which decreased over time (reaching a similar level when compared with the other clusters in the last task periods) (**Figure 5**). Lateral prefrontal cortex is thought to send modulatory signals to sensorimotor areas, by synchronizing at theta-band (Clayton et al. 2015). Our results support an increase of the modulation exerted by FT7 (electrode located in the left lateral prefrontal cortex) over Oz (electrode located in the visual region) in these subjects, suggesting a higher mobilization of attentional resources in the first period of the task (Clayton et al. 2015). However, this modulatory activity reduces over time, indicating that this attentional mobilization decreases progressively over time. Interestingly, it is exactly these subjects that presented a slow-down of RT on the time course of the task. Also regarding point (2) of the model, gamma oscillations in sensory cortices are associated with an enhanced attention to sensory input. Furthermore, the gamma activity in sensory cortices is highly influenced by cognitive control regions (Clayton et al., 2015, Gregoriou et al., 2014). For example, the task improvement of subjects during an oddball task has been associated with the increase of gamma oscillations in visual area (Akimoto et al. 2013; Potes et al. 2014). Interestingly, regarding the increased functional connectivity between FT7 and Oz electrodes, in the present study we also observed an increased (**Figure 6C**), but over-time reduction (**Figure 6D**) of theta-gamma coupling in Oz electrode on subjects included in cluster 2, suggesting an over-time reduced treatment of visual inputs (Gregoriou et al. 2014; Clayton et al. 2015). Altogether, our results highlight that subjects from cluster 2, which are the only ones to have a significant RT slow-down over the time course of the task, showed an increase of functional connectivity between FT7 and Oz electrodes concomitantly with an increase in theta-gamma coupling. In addition, both features reduced over the time course of the task. This suggests that subjects from this cluster have a higher attentional mobilization and increased treatment of visual inputs at the beginning of the task, which decreases over time. Both measures might be used as attentional resources mobilization.

Noteworthy, we observed that subjects from the Cluster 1 showed a widespread increase and over-time maintaining of theta-gamma coupling during the whole task (**Figure 6A** and **Figure 6B**). This may suggest that “high-performers” use a spread brain circuitry lastingly. However, more studies and different techniques are necessary to better explore this feature.

In summary, we were able to identify two populations of individuals presenting opposite speed-accuracy trade-off profiles. We also observed that high-performance subjects present a globally increased theta-gamma coupling maintained over time and an increase of theta activity in FC6 and F8. Finally, we observed that RT slow-down over time, which might indicate the adoption of a more cautious strategy, was observed in individuals showing a high level of modulation exerted by FT7 over Oz at the beginning of the task which strongly reduced during the time course of the task. In addition, a similar result was observed when we looked the theta-gamma coupling in the visual cortex in these subjects, suggesting that both measures may be used as markers of a more controlled strategy. Therefore, the main finding of the present work is that we have been able to identify four attentional profiles by separating subjects based on their performance. This method appears to be very useful in studying and better understanding the mechanisms underlying different attentional features, such as attentional decline, attentional lapses, or speed-accuracy trade-off.

## Compliance with Ethical Standards

### Ethical approval

The study protocol was approved by the local Ethics Committee (“Comité de Protection des Personnes” - CPP Est II).

