## Supplemental data for "Performance - Based Clustering Enables the Study of Physiological Features Supporting Sustained Attention Capacities"

INSERM U1114, Pôle de Psychiatrie, Hôpital Civil de Strasbourg, 1 place de l’Hôpital, 67091 Strasbourg Cedex, France

**SUPPLEMENTARY DATA**

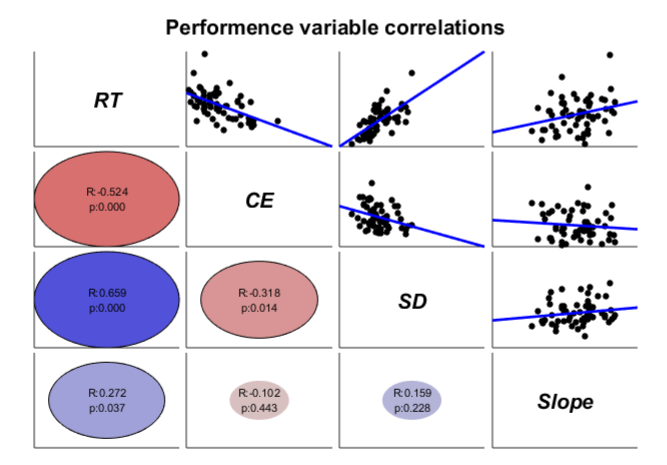

Figure S1: Correlations between performance variables

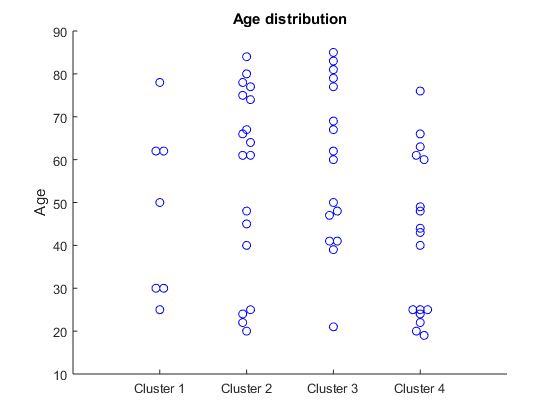

Figure S2: Age distribution among clusters

**Theta power**

**Table S1** shows the level of theta power on monitoring and control processes. We evaluated theta power on the electrodes Fz, AFz, FC5, FC6, FT7, FT8, F7 and F8 electrodes.

| **Table S1:** Theta power. | | | |
| --- | --- | --- | --- |
| **Elect** | **“Cluster”** | **“Period”** | **“Period” Vs “Cluster”** |
| **Fz** | F_(3,275)_= 1.391;  p= 0.255;  cor-p=0.905;  η^2^= 0.07;  power=0.34 | F_(5,275)_= 3.243;  p=0.007;  cor-p=0.054;  η^2^= 0.05;  power=0.88 | F_(15,275)_= 1.391;  p= 0.255;  cor-p=0.905;  η^2^= 0.07;  power=0.34 |
| **AFz** | F_(3,275)_= 2.006;  p= 0.123;  cor-p=0.650;  η^2^= 0.09;  power=0.48 | F_(5,275)_= 3.058;  p=0.010;  cor-p=0.077;  η^2^= 0.05;  power=0.86 | F_(15,275)_= 2.006;  p= 0.123;  cor-p=0.650;  η^2^= 0.09;  power=0.48 |
| **FC5** | F_(3,275)_= 1.782;  p= 0.161;  cor-p=0.754;  η^2^= 0.08;  power=0.48 | F_(5,275)_= 8.670;  p= 0.001;  cor-p=0.007*;  η^2^= 0.13;  power=0.99 | F_(15,275)_= 1.472;  p= 0.114;  cor-p=0.620;  η^2^= 0.07;  power=0.85 |
| **FC6** | F_(3,275)_= 2.250;  p= 0.092;  cor-p=0.537;  η^2^= 0.10;  power=0.53 | F_(5,275)_= 9.664;  p= 0.001;  cor-p=0.007*;  η^2^= 0.14;  power=0.99 | F_(15,275)_= 2.325;  p= 0.003;  cor-p=0.023*;  η^2^= 0.11;  power=0.98 |
| **FT7** | F_(3,275)_= 1.457;  p= 0.236;  cor-p=0.883;  η^2^= 0.07;  power=0.36 | F_(5,275)_= 2.978;  p= 0.012;  cor-p=0.092;  η^2^= 0.05;  power=0.81 | F_(15,275)_= 0.775;  p= 0.704;  cor-p=0.999;  η^2^= 0.04;  power=0.51 |
| **FT8** | F_(3,275)_= 0.745;  p= 0.529;  cor-p=0.997;  η^2^= 0.03;  power=0.19 | F_(5,275)_= 0.571;  p= 0.721;  cor-p=0.999;  η^2^= 0.01;  power=0.20 | F_(15,275)_= 0.748;  p= 0.733;  cor-p=0.999;  η^2^= 0.03;  power=0.49 |
| **F7** | F_(3,275)_= 1.549;  p= 0.212;  cor-p=0.851;  η^2^= 0.07;  power=0.38 | F_(5,275)_= 0.371;  p= 0.868;  cor-p=0.999;  η^2^= 0.006;  power=0.14 | F_(15,275)_= 0.733;  p= 0.749;  cor-p=0.999;  η^2^= 0.03;  power=0.48 |
| **F8** | F_(3,275)_= 2.179;  p= 0.100;  cor-p=0.569;  η^2^= 0.10;  power=0.52 | F_(5,275)_= 7.478;  p= 0.001;  cor-p=0.007*;  η^2^= 0.11;  power=0.99 | F_(15,275)_= 2.232;  p= 0.005;  cor-p=0.039*;  η^2^= 0.10;  power=0.97 |

Regarding the effect of the factor “Period” at **AFz,** we found a higher theta power in period 6 than in periods 2 and 3; at **Fz,** a higher theta power in periods 5 and 6 than in period 2; at **FC5** we found a higher theta power in periods 4, 5 and 6 than in period 2 and in period 6 than in period 1; at **FC6** we found a higher theta power in periods 5 and 6 in comparison to periods 1 and 2; at **FT7** we found a higher theta power in period 5 than in period 1; at F8 we found a higher theta power in periods 5 and 6 than in period 2, but also a higher theta power in period 6 than in period 3. No difference was observed at electrodes **FT8** and **F7**.

The interaction between “Period” and “Cluster” was significant at FC6 and F8 only. In cluster 1, theta power is higher at FC6 in periods 5 and 6 than in periods 1 and 2; and higher at F8 in periods 5 and 6 than in periods 1 and 2.

**Monitoring** – The monitoring activity was measured by looking the modulation in the theta band at Fz and AFz over the electrodes FC5, FC6, FT7, FT8, F7 and F8. No difference was observed over the six task periods. The p-value were corrected for multiple comparisons (cor-p).

| **Table S2:** Cluster effect on monitoring. | | |
| --- | --- | --- |
| Cluster | **F**z | **AF**z |
| **FC5** | F_(3,275)_= 1.292; p=0.286; cor-p = 0.982; η^2^= 0.06; power=0.36 | F_(3,55)_= 0.923; p=0.435; cor-p =0.999; η^2^= 0.04; power=0.23 |
| **FC6** | F_(3,275)_= 0.496; p=0.685; cor-p =0.999; η^2^= 0.02; power=0.14 | F_(3,55)_= 0.306; p=0.820; cor-p =0.999; η^2^= 0.01; power=0.10 |
| **FT7** | F_(3,275)_= 1.479; p=0.230; cor-p =0.956; η^2^= 0.07; power=0.36 | F_(3,55)_= 1.097; p=0.357; cor-p =0.995; η^2^= 0.05; power=0.28 |
| **FT8** | F_(3,275)_= 0.902; p=0.445; cor-p =0.999; η^2^= 0.04; power=0.23 | F_(3,55)_= 0.361; p=0.780; cor-p =0.999; η^2^= 0.01; power=0.11 |
| **F7** | F_(3,275)_= 2.462; p=0.072; cor-p =0.592; η^2^= 0.11; power=0.58 | F_(3,55)_= 1.340; p=0.270; cor-p =0.977; η^2^= 0.06; power=0.33 |
| **F8** | F_(3,275)_= 1.747; p=0.168; cor-p =0.889; η^2^= 0.08; power=0.43 | F_(3,55)_= 1.390; p=0.255; cor-p =0.970; η^2^= 0.07; power=0.34 |

| **Table S3:** Period effect on monitoring. | | |
| --- | --- | --- |
| Period | **F**z | **AF**z |
| **FC5** | F_(5,275)_= 1.284; p=0.270; cor-p =0.977; η^2^= 0.02; power=0.45 | F_(5,55)_= 0.718; p=0.609; cor-p =0.999; η^2^= 0.01; power=0.25 |
| **FC6** | F_(5,275)_= 1.320; p=0.255; cor-p =0.970; η^2^= 0.02; power=0.46 | F_(5,55)_= 1.040; p=0.394; cor-p =0.997; η^2^= 0.01; power=0.36 |
| **FT7** | F_(5,275)_= 0.154; p=0.978; cor-p =1; η^2^< 0.00; power=0.08 | F_(5,55)_= 1.300; p=0.263; cor-p =0.974; η^2^= 0.02; power=0.45 |
| **FT8** | F_(5,275)_= 0.800; p=0.550; cor-p =0.999; η^2^= 0.01; power=0.28 | F_(5,55)_= 0.676; p=0.641; cor-p =0.999; η^2^= 0.01; power=0.24 |
| **F7** | F_(5,275)_= 0.320; p=0.900; cor-p =1; η^2^< 0.00; power=0.13 | F_(5,55)_= 0.581; p=0.713; cor-p =0.999; η^2^= 0.01; power=0.22 |
| **F8** | F_(5,275)_= 0.964; p=0.440; cor-p =0.999; η^2^= 0.01; power=0.34 | F_(5,55)_= 0.531; p=0.752; cor-p =0.999; η^2^< 0.00; power=0.19 |

| **Table S4:** Interaction between cluster and period on monitoring. | | |
| --- | --- | --- |
| Interaction | **F**z | **AF**z |
| **FC5** | F_(15,275)_= 1.338; p=0.178; cor-p =0.904; η^2^= 0.06; power=0.81 | F_(15,275)_= 1.905; p=0.022; cor-p =0.234; η^2^= 0.09; power=0.94 |
| **FC6** | F_(15,275)_= 0.767; p=0.713; cor-p =0.999; η^2^= 0.04; power=0.50 | F_(15,275)_= 1.203; p=0.268; cor-p =0.976; η^2^= 0.06; power=0.75 |
| **FT7** | F_(15,275)_= 0.569; p=0.897; cor-p =0.999; η^2^= 0.03; power=0.37 | F_(15,275)_= 0.449; p=0.962; cor-p =1; η^2^= 0.02; power=0.28 |
| **FT8** | F_(15,275)_= 1.279; p=0.214; cor-p =0.944; η^2^= 0.06; power=0.78 | F_(15,275)_= 1.014; p=0.440; cor-p =0.999; η^2^= 0.05; power=0.65 |
| **F7** | F_(15,275)_= 0.675; p=0.807; cor-p =0.999; η^2^= 0.03; power=0.44 | F_(15,275)_= 0.583; p=0.886; cor-p =1; η^2^= 0.03; power=0.38 |
| **F8** | F_(15,275)_= 0.844; p=0.067; cor-p =0.564; η^2^= 0.04; power=0.55 | F_(15,275)_= 1.322; p=0.187; cor-p =0.916; η^2^= 0.06; power=0.80 |

Only the Granger connectivity between AFz and FC5 showed an interaction between the factors “Period” and “Cluster” (**Table S4**). In Cluster 1, theta power is lower in periods 5 and 6 than in period 2; in Cluster 4, theta power is lower in periods 1 and 6 than in period 5 (**Figure S1**).

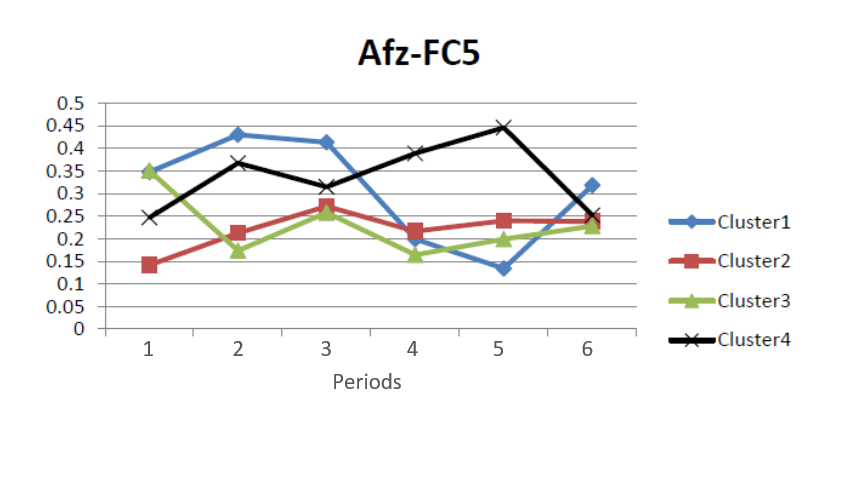
**Figure S3**: Connectivity between AFz and FC5.

**AFz-FC5**

**Activation of task-relevant regions** - The modulation of task-relevant regions was evaluated by looking at the coherence at theta band at electrodes FC5, CF6, FT7, FT8, F7 and F8 and electrodes located in visual regions (Oz, POz, PO3 and PO4). The p-values were corrected for multiple comparisons (cor-p).

| **Table S5:** Cluster effect on activation of task-relevant regions. | | | | | | |
| --- | --- | --- | --- | --- | --- | --- |
| **Cluster** | **FC5** | **FC6** | **FT7** | **FT8** | **F7** | **F8** |
| **O**z | F_(3,55)_= 1.208; p=0.315;  cor-p= 0.999;  η^2^= 0.06; power=0.30 | F_(3,55)_= 2.079; p=0.113;  cor-p= 0.943;  η^2^= 0.10; power=0.50 | F_(3,55)_= 1.439; p=0.241;  cor-p= 0.998;  η^2^= 0.07; power=0.36 | F_(3,55)_= 0.792; p=0.503;  cor-p= 0.999;  η^2^= 0.04; power=0.20 | F_(3,55)_= 0.532; p=0.662;  cor-p= 1;  η^2^= 0.02; power=0.15 | F_(3,55)_= 0.848; p=0.473;  cor-p= 0.999;  η^2^= 0.04; power=0.22 |
| **PO**z | F_(3,55)_= 1.416; p=0.247;  cor-p= 0.998;  η^2^= 0.07; power=0.35 | F_(3,55)_= 1.733; p=0.170;  cor-p= 0.998;  η^2^= 0.08; power=0.42 | F_(3,55)_= 1.110; p=0.352;  cor-p= 0.999;  η^2^= 0.05; power=0.28 | F_(3,55)_= 0.929; p=0.432;  cor-p= 0.999;  η^2^= 0.04; power=0.24 | F_(3,55)_= 0.906; p=0.443;  cor-p= 0.999;  η^2^= 0.04; power=0.23 | F_(3,55)_= 1.211; p=0.314;  cor-p= 0.999;  η^2^= 0.06; power=0.30 |
| **PO3** | F_(3,55)_= 2.267; p=0.090;  cor-p= 0.896;  η^2^= 0.11; power=0.54 | F_(3,55)_= 2.459; p=0.072;  cor-p= 0.833;  η^2^= 0.11; power=0.58 | F_(3,55)_= 4.663; p=0.005;  cor-p= 0.113;  η^2^= 0.20; power=0.87 | F_(3,55)_= 1.713; p=0.174;  cor-p= 0.989;  η^2^= 0.08; power=0.42 | F_(3,55)_= 1.684; p=0.180;  cor-p= 0.991;  η^2^= 0.08; power=0.41 | F_(3,55)_= 2.533; p=0.066;  cor-p= 0.805;  η^2^= 0.12; power=0.59 |
| **PO4** | F_(3,55)_= 0.553; p=0.647;  cor-p= 1;  η^2^= 0.02; power=0.15 | F_(3,55)_= 1.177; p=0.326;  cor-p= 0.999;  η^2^= 0.06; power=0.29 | F_(3,55)_= 0.784; p=0.507;  cor-p= 0.999;  η^2^= 0.04; power=0.20 | F_(3,55)_= 0.824; p=0.486;  cor-p= 0.999  η^2^= 0.04; power=0.21 | F_(3,55)_= 0.473; p=0.702;  cor-p= 1;  η^2^= 0.02; power=0.13 | F_(3,55)_= 0.419; p=0.740;  cor-p= 1;  η^2^= 0.02; power=0.12 |

An effect of the factor cluster was observed between FT7-PO3 (**Table S5**) due to a higher correlation in the cluster 2 compared to the clusters 3 and 4.

| **Table S6:** Period effect on activation of task-relevant regions. | | | | | | |
| --- | --- | --- | --- | --- | --- | --- |
| **Period** | **FC5** | **FC6** | **FT7** | **FT8** | **F7** | **F8** |
| **O**z | F_(5,275)_= 1.344; p=0.245;  cor-p= 0.998;  η^2^= 0.02; power=0.47 | F_(5,275)_= 3.497; p=0.004;  cor-p= 0.091;  η^2^= 0.05; power=0.91 | F_(5,275)_= 0.756; p=0.581;  cor-p= 0.999;  η^2^= 0.01; power=0.27 | F_(5,275)_= 0.542; p=0.744;  cor-p= 1;  η^2^< 0.00; power=0.19 | F_(5,275)_= 0.574; p=0.719;  cor-p= 1;  η^2^= 0.01; power=0.21 | F_(5,275)_= 0.348; p=0.882;  cor-p= 1;  η^2^< 0.00; power=0.13 |
| **PO**z | F_(5,275)_= 0.190; p=0.965;  cor-p= 1;  η^2^< 0.00; power=0.35 | F_(5,275)_= 0.996; p=0.419;  cor-p= 0.999;  η^2^= 0.01; power=0.35 | F_(5,275)_= 0.533; p=0.750;  cor-p= 1;  η^2^< 0.00; power=0.19 | F_(5,275)_= 1.348; p=0.244;  cor-p= 0.998;  η^2^= 0.02; power=0.47 | F_(5,275)_= 2.244; p=0.050;  cor-p= 0.708;  η^2^= 0.03; power=0.72 | F_(5,275)_= 0.374; p=0.865;  cor-p= 1;  η^2^< 0.00; power=0.14 |
| **PO3** | F_(5,275)_= 1.367; p=0.236;  cor-p= 0.998;  η^2^= 0.02; power=0.48 | F_(5,275)_= 1.033; p=0.398;  cor-p= 0.999;  η^2^= 0.01; power=0.36 | F_(5,275)_= 1.974; p=0.082;  cor-p= 0.871;  η^2^= 0.03; power=0.66 | F_(5,275)_= 0.549; p=0.738;  cor-p= 1;  η^2^< 0.00; power=0.20 | F_(5,275)_= 0.807; p=0.544;  cor-p= 0.999;  η^2^= 0.01; power=0.28 | F_(5,275)_= 0.316; p=0.902;  cor-p= 1;  η^2^< 0.00; power=0.12 |
| **PO4** | F_(5,275)_= 1.063; p=0.380;  cor-p= 0.999;  η^2^= 0.01; power=0.37 | F_(5,275)_= 2.049; p=0.072;  cor-p= 0.833;  η^2^= 0.03; power=0.67 | F_(5,275)_= 1.905; p=0.093;  cor-p= 0.903;  η^2^= 0.03; power=0.64 | F_(5,275)_= 0.886; p=0.490;  cor-p= 0.999;  η^2^= 0.01; power=0.31 | F_(5,275)_= 1.103; p=0.358;  cor-p= 0.999;  η^2^= 0.01; power=0.39 | F_(5,275)_= 0.665; p=0.650;  cor-p= 0.999;  η^2^= 0.01; power=0.24 |

An effect of the factor period was found for the comparison between FC6-Oz (**Table S6**) due to a higher correlation of period 1 compared to periods 3, 5 and 6.

| **Table S7:** Interaction between cluster and period on activation of task-relevant regions. | | | | | | |
| --- | --- | --- | --- | --- | --- | --- |
| **Interaction** | **FC5** | **FC6** | **FT7** | **FT8** | **F7** | **F8** |
| **O**z | F_(15,275)_= 1.457; p=0.120;  cor-p= 0.953;  η^2^= 0.07; power=0.85 | F_(15,275)_= 1.882; p=0.024;  cor-p= 0.441;  η^2^= 0.09; power=0.94 | F_(15,275)_= 2.628; p<0.001;  cor-p= 0.023;*  η^2^= 0.12; power=0.99 | F_(15,275)_= 1.644; p=0.062;  cor-p= 0.784;  η^2^= 0.08; power=0.90 | F_(15,275)_= 0.907; p=0.556;  cor-p= 0.999;  η^2^= 0.04; power=0.59 | F_(15,275)_= 0.992; p=0.463;  cor-p= 0.999;  η^2^= 0.05; power=0.64 |
| **PO**z | F_(15,275)_= 0.819; p=0.655;  cor-p= 1;  η^2^= 0.04; power=0.54 | F_(15,275)_= 1.163; p=0.300;  cor-p= 0.999;  η^2^= 0.05; power=0.73 | F_(15,275)_= 1.396; p=0.148;  cor-p= 0.978;  η^2^= 0.07; power=0.83 | F_(15,275)_= 2.192; p=0.006;  cor-p= 0.134;  η^2^= 0.10; power=0.97 | F_(15,275)_= 0.578; p=0.890;  cor-p= 1;  η^2^= 0.03; power=0.37 | F_(15,275)_= 0.523; p=0.926;  cor-p= 1;  η^2^= 0.02; power=0.33 |
| **PO3** | F_(15,275)_= 1.771; p=0.038;  cor-p= 0.605;  η^2^= 0.08; power=0.92 | F_(15,275)_= 1.024; p=0.429;  cor-p= 0.999  η^2^= 0.05; power=0.66 | F_(15,275)_= 2.852; p<0.001;  cor-p= 0.023;*  η^2^= 0.13; power=0.99 | F_(15,275)_= 1.547; p=0.088;  cor-p= 0.890;  η^2^= 0.07; power=0.87 | F_(15,275)_= 0.638; p=0.842;  cor-p= 1;  η^2^= 0.03; power=0.41 | F_(15,275)_= 0.540; p=0.916;  cor-p= 1;  η^2^= 0.02; power=0.35 |
| **PO4** | F_(15,275)_= 0.894; p=0.570;  cor-p= 0.999;  η^2^= 0.04; power=0.58 | F_(15,275)_= 1.655; p=0.059;  cor-p= 0.767;  η^2^= 0.08; power=0.90 | F_(15,275)_= 1.347; p=0.173;  cor-p= 0.989;  η^2^= 0.06; power=0.81 | F_(15,275)_= 1.979; p=0.016;  cor-p= 0.320;  η^2^= 0.09; power=0.95 | F_(15,275)_= 1.071; p=0.382;  cor-p= 0.999;  η^2^= 0.05; power=0.69 | F_(15,275)_= 0.902; p=0.561;  cor-p= 0.999;  η^2^= 0.04; power=0.59 |

The ANOVA of Granger connectivity showed an interaction between “Period” and “Cluster” for the connection between FC5-PO3 due to a higher theta activity on the period1 compared to the periods 3, 4 and 6 in the cluster 2; FC6-Oz due to a higher theta activity on the period 1 compared to the periods 5 and 6 in the cluster 2; FT7-Oz due to a higher theta activity on the period 1 compared to the periods 5 and 6 in the cluster 2; FT7-PO3 due to a higher theta activity on the period 1, 2, 3 and 4 compared to the period 6 and a higher theta activity in the period 1 compared to the period 5 in the cluster 2; FT8-POz due to a higher theta activity on the period 6 compared to the periods 1 and 2 in the cluster 4 and FT8-PO4 due to a higher theta activity on the period 4 compared to the period 2 in the cluster 4 (**Table S7; Figure S2**).

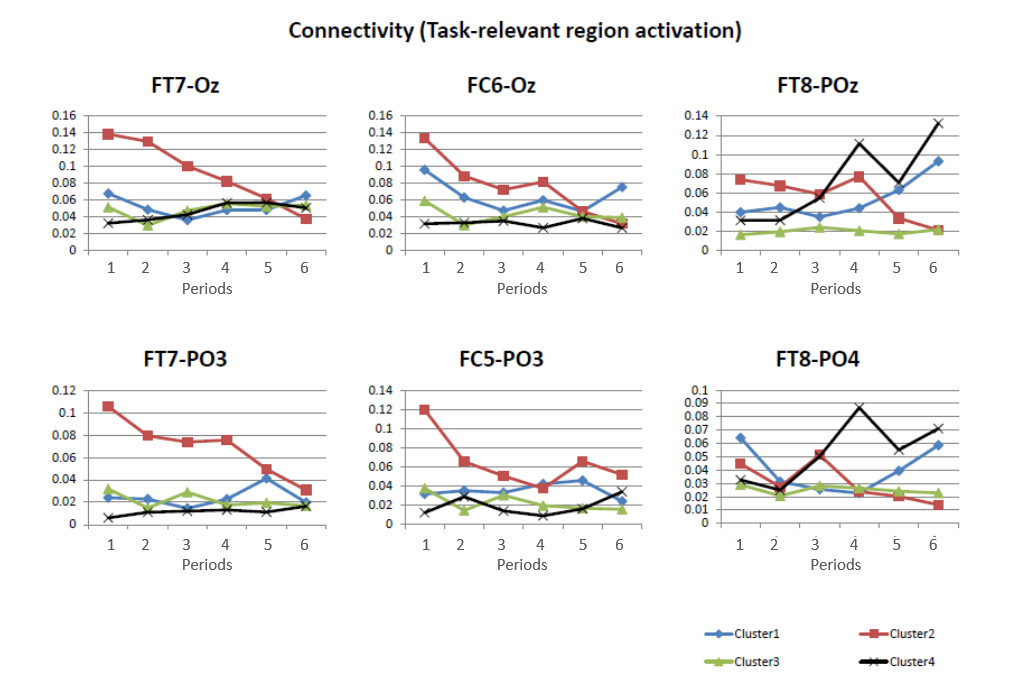
**Figure S4**: Connectivity between monitoring-related and task-relevant regions.
